# The effects of haploid selection on Y chromosome evolution in two closely related dioecious plants

**DOI:** 10.1101/264382

**Authors:** George Sandler, Felix E.G. Beaudry, Spencer C.H. Barrett, Stephen I. Wright

## Abstract

The evolution of sex chromosomes is usually considered to be driven by sexually antagonistic selection in the diploid phase. However, selection during the haploid gametic phase of the lifecycle has recently received theoretical attention as possibly playing a central role in sex chromosome evolution, especially in plants where gene expression in the haploid phase is extensive. In particular, male-specific haploid selection might favour the linkage of pollen beneficial alleles to male sex determining regions on incipient Y chromosomes. This linkage might then allow such alleles to further specialise for the haploid phase. Purifying haploid selection is also expected to slow the degeneration of Y-linked genes expressed in the haploid phase. Here, we examine the evolution of gene expression in flower buds and pollen of two species of *Rumex* to test for signatures of haploid selection acting during plant sex chromosome evolution. We find that genes with high ancestral pollen expression bias occur more often on sex chromosomes than autosomes and that genes on the Y chromosome are more likely to become enriched for pollen expression bias. We also find that genes with low expression in pollen are more likely to be lost from the Y chromosome. Our results suggest that sex-specific haploid selection during the gametophytic stage of the lifecycle may be a major contributor to several features of plant sex chromosome evolution.

## IMPACT SUMMARY

Selection in the haploid phase of the life cycle is considered to be a strong force allowing for the efficient purging and fixation of recessive alleles. Previous theoretical and empirical work suggests that haploid selection can affect plant sex chromosome evolution in several ways. Haploid selection should favour the suppression of recombination allowing haploid beneficial alleles to fix on chromosomes that segregate into the sex experiencing stronger haploid selection, generally males. Haploid selection may also allow such genes to subsequently specialise for this haploid stage. Finally, purifying haploid selection may slow down the degeneration of non-recombining chromosomes. Evidence for these processes is, however, limited. Here, we analyse gene expression data from three tissues of two *Rumex* species to look for signals of haploid selection acting on plant Y chromosome evolution. We demonstrate that the sex chromosomes in these species are enriched for pollen-expressed genes, that the genes have become more pollen biased in expression, and that Y-linked genes are overexpressed in pollen. Our results support previous findings in *Silene* that haploid selection contributes to the retention of genes on the Y chromosome, but also provides novel empirical evidence for adaptive specialization of Y-linked genes for the haploid phase of the plant life cycle.

Theory suggests sex chromosomes evolve from a pair of autosomes that acquire a sex-determining region and subsequently accumulate sexually antagonistic alleles expressed in the diploid phase. The loss of genetic recombination between the sex chromosomes is thought to be selected to assure the segregation of sexually antagonistic alleles into the sex in which they are beneficial (Rice 1984; Lenormand 2003). Though widely accepted, evidence supporting sex-specific selection is limited to a few systems (Foerster et al. 2007; Delph et al. 2010; Innocenti and Morrow 2010) and has rarely been conclusively related to sex chromosome evolution (but see Wright et al. 2017).

A key feature of the angiosperm life-cycle is the predominance of a haploid gametophytic phase (Haldane 1933). In the hermaphroditic plant *Arabidopsis thaliana* 60-70% of all genes are expressed in pollen (Honys and Twell 2004; Borges et al. 2008). Such haploid expression exposes genes to a unique selective regime which includes more efficient removal of deleterious mutations from a population as recessive deleterious phenotypic effects are expressed (Gerstein and Otto 2009). Similarly, recessive beneficial mutations are more likely to spread through populations. Indeed, in *Capsella grandiflora*, pollen-expressed genes experience stronger purifying and positive selection relative to non-pollen expressed genes (Arunkumar et al. 2013). Pollen competition is generally considered to be a common feature of angiosperms, further increasing selective pressures imposed on plant genomes in the haploid phase (Moore and Pannell 2011).

Gene expression in pollen may contribute to the evolution of sex chromosomes in three complementary ways (Fig. 1). First, similar to sexually antagonistic alleles in diploids, pollen-specific (and therefore sex-specific) haploid selection can favour the loss of recombination between the X and Y chromosomes, because linkage of alleles beneficial during pollen competition to the Y chromosome enables these alleles to spend more time in males where competition occurs (Scott and Otto 2017). Hereafter, we refer to this phenomenon as adaptive linkage (Fig. 1a). Second, once genes have become sex-linked, greater haploid selection in males may cause divergence and upregulation of Y-linked alleles specialized for pollen (hereafter “pollenization”), akin to masculinization of the Y (Lahn and Page 1997; Zhou and Bachtrog 2012) and feminization of the X (Prince et al. 2010; Allen et al. 2013; Albritton et al. 2014) observed in some animal systems (Fig. 1b). Finally, haploid expression of genes on the Y can cause biased retention of pollen-expressed genes from the degenerating Y, as reported in *Silene latifolia* (Chibalina and Filatov 2011) (Fig. 2c).The study of pollen expression in young plant sex chromosome systems provides opportunities for the untangling of these three processes.

**Fig. 1:**
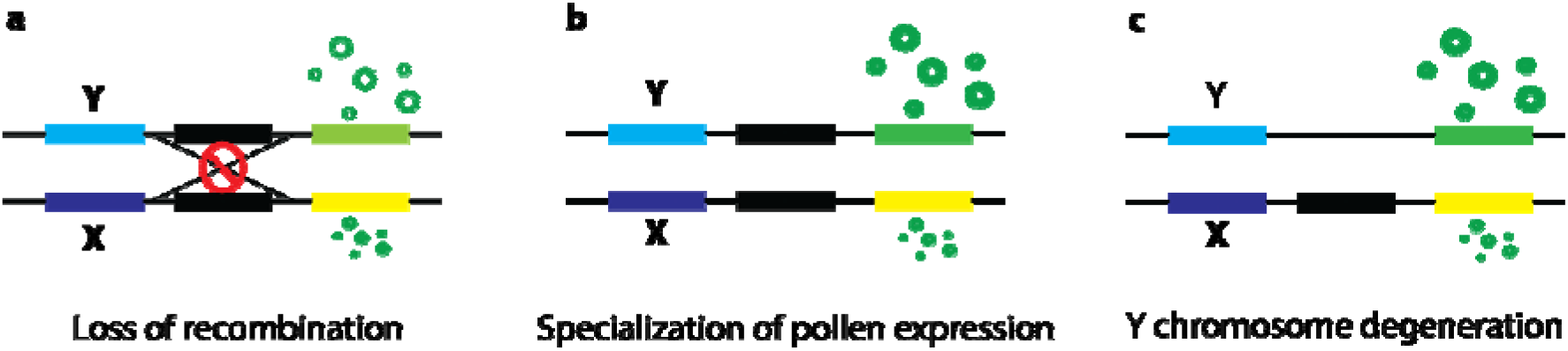
Depiction of the effects of haploid gametophytic selection on sex chromosomes. Three distinct processes can potentially contribute to biased expression and overrepresentation of pollen genes on the Y chromosome. **a)** Recombination can be lost between the male determining region (Y) and any allele that increases pollen fitness (the green allele). **b)** Without recombination, alleles with pollen-specific fitness can diverge, or their expression can increase relative to the X-linked allele. **c)** As inefficient selection due to linkage causes degeneration of Y-linked alleles, haploid selection during pollen competition may cause biased retention of genes with pollen-specific fitness effects

**Fig. 2:**
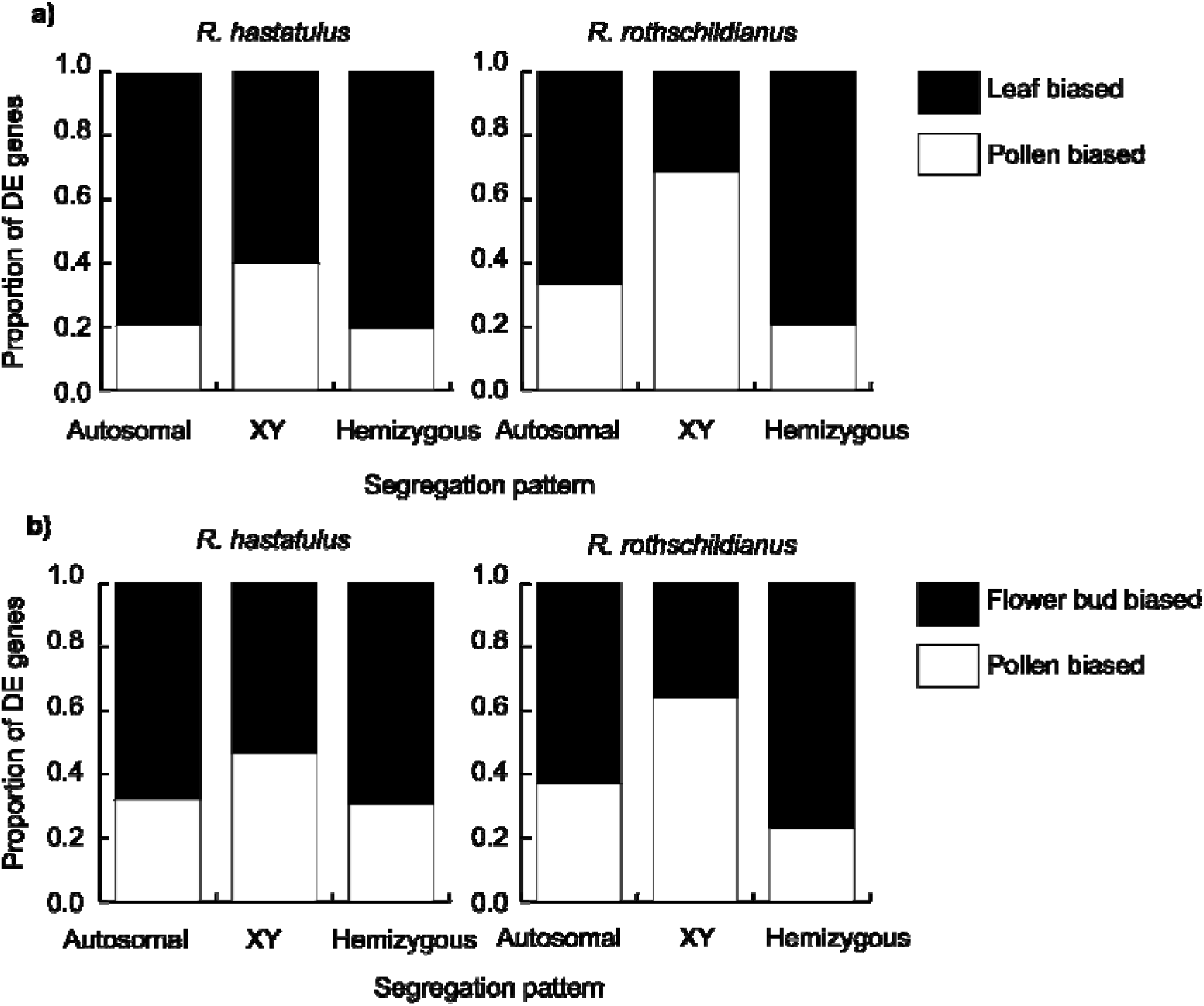
Tissue expression bias of different gene groups in two *Rumex* species. Bar segments represent the fraction of genes with significant differential-expression (DE) in two pairwise tissue comparisons.

*Rumex* (Polygonaceae) provides a valuable study system to investigate the effects of sex-specific haploid selection and sexual antagonism in the diploid phase on sex chromosome evolution. Members of this genus possess uniovulate flowers, and open-pollinated flowers capture numerous pollen grains increasing the scope for pollen competition and haploid selection (Stehlik et al. 2008). Here, we sequence RNA from flower buds and mature pollen of annual, wind-pollinated *R. hastatulus* and *R. rothschildianus* to investigate patterns of gene expression and test the predictions of models of sex chromosome evolution driven by sexually antagonistic and haploid selection. Both species possess heteromorphic sex chromosomes, with non-orthologous sex-linked genes, consistent with independently-evolved sex chromosomes (Crowson et al. 2017). The species also exhibit different degrees of Y-chromosome degeneration (Crowson et al. 2017). In *R. hastatulus* approximately three-quarters of genes have retained an expressed Y copy, whereas in *R. rothschildianus* only ~10% have been retained (Hough et al. 2014; Crowson et al. 2017). To investigate the role of sex-specific haploid selection in the evolution of plant sex chromosomes we sought to address the following questions using our two focal species of *Rumex*: 1) Were the sex chromosomes ancestrally enriched for pollen-biased genes, as expected if the spread of recombination was driven by male haploid selection? 2) Is there evidence of subsequent pollenization of Y-linked genes? 3) Does haploid selection preserve pollen-expressed genes on degenerating Y chromosomes?

## METHODS

### Tissue collection

We used gene expression data from three tissues: pollen (a target for haploid selection), male flower bud (a target for sexual antagonism) and male leaf (control). We collected mature pollen and filtered it through a fine nylon mesh before RNA extraction. We pooled pollen from two *R. rothschildianus* males due to low tissue yields in this species but collected pollen from two male plants of *R. hastatulus* individually. We collected developing, unopened male flower buds from two *R. rothschildianus* individuals (sampled and sequenced independently) and one *R. hastatulus* individual. We performed all RNA extractions using Spectrum™ Plant Total RNA kits and stored RNA at −80°C. Leaf expression data from three *R. hastatulus* and three *R. rothschildianus* males were obtained from previous work (see Hough et al. 2014; Crowson et al. 2017).

### RNAseq and read analysis

We sequenced RNA samples using Illumina Hi-seq 2500 sequencing with 100bp paired end reads at the Centre for Applied Genomics, Toronto. We aligned samples to existing female leaf transcriptome assemblies from both species (Hough et al. 2014; Crowson et al. 2017). We performed alignments using STAR (Dobin et al. 2013) after which we removed duplicate reads using Picard (http://broadinstitute.github.io/picard). We used SAMtools to retrieve read counts for downstream differential expression analysis (Li et al. 2009). We performed differential expression analysis using the R package DESeq2 (Love et al. 2014) using read counts obtained from SAMtools. We used >0.3 FPKM as a cut-off for active transcription, as recommended in (Ramsköld et al. 2009).

### Ortholog comparison

We compared expression of genes that were retained on both the X and Y (hereafter XY genes) and their orthologs in *R. hastatulus* and *R. rothschildianus* to infer changes in expression of genes that had become sex linked since the species diverged. This is possible because the sex-linked genes in the two species of *Rumex* appear to have arisen independently (Crowson et al. 2017). We obtained lists of orthologs between the species from Crowson et al. (2017). Because we were interested in the relative expression bias of XY linked genes, and the normalization of expression may be influenced by the proportion and number of sex-linked genes in the genome, we corrected for this by normalizing expression bias of XY genes by the average expression bias of autosomes. The method for calculating the average autosomal expression bias is given by:

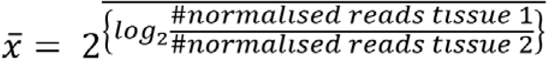

After we calculated the average autosomal expression bias, we corrected individual XY gene bias using equation 2:

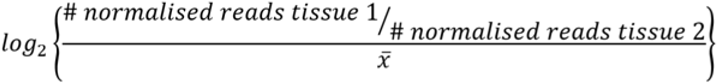

### SNP calling and analysis of allele-specific expression

We used the HaplotypeCaller tool in GATK (McKenna et al. 2010) to call SNP variants in our transcriptomes followed by the SelectVariants tool in GATK to select only biallelic, sex-linked SNPs. This list of SNPs was then run through the GATK ASEReadcounter (McKenna et al. 2010) tool with default settings to allow for estimation of allele-specific expression on the sex chromosomes. As all male tissue transcriptomes were aligned to female references, most alternate SNPs in XY linked genes represent the Y copy of a gene whereas most reference SNPs represent the X copy. However, this assumption can be violated if a polymorphism exists on the X. To account for this, we used population data from *R. hastatulus* to identify fixed SNP differences between the X and Y chromosomes (Hough et al. 2017). Fixed differences were determined if all females in a population were homozygous at an X linked SNP whereas all males were heterozygous for the same site. This yielded a list of high confidence sites for use in allele-specific expression analysis. No such population data was available for *R. rothschildianus* so instead we used SNPs with sex-linked segregation patterns within a family of sequenced plants (Crowson et al. 2017).

The lists of high confidence sites for allele-specific expression analysis were then run through the tool GeneiASE (Edsgärd et al. 2016) to find statistically significant cases of allelic bias in expression. Since GeneiASE does not require phasing, the output identifies which genes have a bias in expression but not the direction of this bias. We averaged the log 2-fold expression change of each high confidence XY site to infer whether the reference (X) or alternate (Y) copy was overexpressed.

### Statistical analysis

All *P*-values reported in this study are two tailed. We used Fisher’s exact tests to identify significant differences between counts of differentially expressed genes. Student’s *t*-tests were used to compare mean FPKM pollen expression of hemizygous and XY linked genes. We conducted Wilcoxon signed-rank tests to compare tissue expression bias of XY and ortholog genes. Multiple test correction was applied by both DESeq2 and GeneiASE using the Benjamin-Hochberg method, only genes with an adjusted *P*-value of <0.05 were considered as differentially expressed (for DESeq2), or as having significant allele-biased expression (for GeneiASE). We used Fisher’s combined probability tests to combine the individual gene *P*-values using GeneiASE for each tissue sample to yield one *P*-value per gene. We then used Fisher’s exact tests to determine differences in the numbers of X vs. Y biased genes in our allele-specific expression analysis. We did not test for differences in numbers of genes without allele-specific expression, because the fraction of non-biased genes heavily depends on factors such as sequencing depth, which can vary from sample to sample. We used counts of overexpressed genes in leaves as the null expected counts in our statistical tests, as we did not expect either sex-specific haploid selection or diploid sexual antagonism to be driving Y chromosome evolution in leaf tissue.

## Result and Discussion

### Gene expression is widespread in pollen

In *R. hastatulus*, 39.1% of all predicted leaf transcripts had signatures of active pollen transcription, whereas 50.9% of leaf transcripts in *R. rothschildianus* showed evidence of pollen expression. These results suggest widespread gene expression in the haploid phase. The values we obtained for pollen were significantly lower than in flower bud tissue, where 81.7% and 87.3% of predicted leaf transcripts were actively transcribed in *R. hastatulus* and *R. rothschildianus*, respectively. Despite overlap between tissues in genes with active expression, principle components analysis (Supplementary Fig. 1) of individual samples indicated strong differentiation between tissues in expression, particularly for pollen.

### Sex chromosomes are enriched for haploid expressed genes

To investigate the relative importance of sexual antagonism and haploid selection during sex chromosome evolution, we compared sex-linked and autosomal genes for expression bias across three tissues. We focused on expression differences between male leaf, male developing flower bud and mature pollen. The chromosomal location of genes was previously evaluated using SNP segregation patterns for both *R. hastatulus* (Hough et al. 2014) and *R. rothschildianus* (Crowson et al. 2017). We compared counts of genes identified as having significantly different expression (Benjamin-Hochberg FDR adjusted *P* < 0.05) between two pairs of tissues to quantify expression bias. When comparing leaf and pollen expression patterns (Fig. 2a), we found that sex-linked genes with retained Y copies (hereafter XY genes) were more often significantly pollen biased than autosomal genes in both *R. hastatulus* (Fisher’s exact test, *P* < 0.0001) and *R. rothschildianus* (Fisher’s exact test, *P*< 0.0001). The same was true when comparing pollen to flower bud expression in *R. hastatulus* (Fisher’s exact test, *P* = 0.0006) and *R. rothschildianus* (Fisher’s exact test, *P* < 0.0001) (Fig. 2b). However, this enrichment of pollen-biased genes on the sex chromosomes could be driven by three distinct factors (Fig. 1); an ancestral bias due to selection for sex-linkage, pollenization following the formation of the sex chromosomes, and differential degeneration due to haploid expression. Our subsequent analyses sought to evaluate the relative contribution of these various processes.

### Haploid selection maintains pollen-expressed genes on the Y chromosome

To test whether purifying selection and haploid expression of Y-linked genes slow down Y chromosome degeneration in *Rumex* (Fig. 1c), we compared the pollen expression of hemizygous genes (which lack a Y-expressed copy) and XY genes (Chibalina and Filatov 2011; Crowson et al. 2017). We found that hemizygous genes showed significantly reduced pollen expression compared with XY genes in *R. hastatulus* (Welch Two Sample *t*-test, *t* = −6.7295, df = 154.31, *P* = 3.145e-10) and *R. rothschildianus* (Welch Two Sample *t*-test, *t* = −14.019, df = 552.18, *P* = 2.2e-16) (Supplementary Fig. 2). This effect is particularly prominent in the more degenerated *R. rothschildianus:* the effect size (Cohen’s D) in *R. hastatulus* is 0.74, whereas it is 1.16 in *R. rothschildianus*. This difference was not simply due to hemizygous genes being generally less expressed (Crowson et al. 2017); indeed, differential expression analyses indicated that hemizygous genes had a deficiency of pollen-biased relative to either leaf- or flower-biased genes compared with XY genes in both *R. hastatulus* (Fisher’s exact test, *P* < 0.0001 leaf/pollen; *P* < 0.0252 flower bud/pollen) and *R. rothschildianus* (Fisher’s exact test, *P* < 0.0001 leaf/pollen; *P* < 0.0001 flower bud/pollen) (Fig. 2). Again, this effect was more pronounced in *R. rothschildianus*. Our results suggest that haploid selection retains pollen expressed genes, as also reported in *Silene latifolia*, another XY plant sex chromosome system (Chibalina and Filatov 2011). It is interesting to note that the difference in pollen expression between XY and hemizygous genes has diverged to a greater extent in *R. rothschildianus* suggesting that haploid selection does indeed slow down Y chromosome degeneration even in a highly heteromorphic plant sex chromosome system.

### Sex chromosome linked genes show signals of ancestral and derived pollen bias

We next investigated whether the footprint of differential Y chromosome degeneration could fully account for the patterns of pollen bias at XY genes, without needing to invoke adaptive evolution of sex linkage or Y chromosome pollenization. To do this, we examined the extent of pollen bias on all sex chromosome-linked genes, combining both hemizygous and XY genes. By combining these gene sets, our analysis should more closely resemble the ancestral set of genes that evolved to become linked to the sex-determining region prior to Y chromosome degeneration. The combined data still showed an enrichment of pollen-biased genes across all sex-linked genes analysed together for both *R. hastatulus* (Fisher’s exact test, *P* < 0.0001, *P* = 0.0015; pollen/leaf and pollen/flower bud respectively) and *R. rothschildianus* (Fisher’s exact test, *P* = 0.0005, *P* = 0.0314; pollen/leaf and pollen/flower bud, respectively) (Supplementary Fig. 3). However, there exists an ascertainment bias in this reconstructed gene set as more SNP segregation patterns can be used to identify XY genes compared to hemizygous genes (in particular, divergent SNPs between X and Y chromosomes) resulting in overrepresentation of XY genes on reconstructed XY chromosomes (Hough et al. 2014; Crowson et al. 2017). To account for this bias, we used existing published XY gene lists (Hough et al. 2014; Crowson et al. 2017) which only contained XY genes identified with the same set of SNP segregation patterns (polymorphisms on the X chromosome) as hemizygous genes. This procedure therefore removed any ascertainment bias. We still found evidence for a significant enrichment of pollen-biased genes in *R. hastatulus* (Fisher’s exact test, *P* < 0.0001 for both tissue comparisons), but not in *R. rothschildianus*, where sex-linked genes as a whole were significantly depleted for pollen enrichment (Fisher’s exact test, *P* = 0.0224, *P* = 0.0074; pollen/leaf and pollen/flower bud, respectively). Thus, overall the enrichment of pollen-expressed genes does not appear to be a simple function of the Y degeneration of genes not expressed in pollen. However, it is possible that any signal of early enrichment may have been eroded by extensive Y degeneration in the more degenerated *R. rothschildianus* and possibly secondary movement of genes on and off the X chromosome.

Because Y-chromosome degeneration alone is not sufficient to explain the enrichment of pollen expression on XY linked genes, we investigated whether adaptive linkage (Fig. 1a), and/or pollenization (Fig. 1b) of the Y chromosome could account for differential pollen expression between sex-linked and autosomal genes. The sex-linked genes in *R. rothschildianus* and *R. hastatulus* have arisen independently (Crowson et al. 2017), thus ancestral expression of XY linked genes in one species should be represented by the autosomal orthologs of these genes in the other species. Therefore, we next attempted to disentangle the ancestral and subsequent evolution of expression bias in sex-linked genes.

We first investigated whether orthologs of XY-linked genes were ancestrally more pollen biased than other autosomal genes to determine whether pollen bias is present before linkage to the sex chromosomes, which would be indicative of adaptive sex linkage. We found that *R. hastatulus* XY-linked genes were ancestrally more pollen biased than other autosomal genes in a comparison between leaf and pollen (Fisher’s exact test, *P* = 0.0344) (Supplementary Fig. 4); although a similar trend was evident comparing pollen and flower buds, the difference was not significant (Fisher’s exact test, *P* = 0.1197). Similarly, *R. rothschildianus* XY-linked genes were ancestrally more pollen biased than other autosomal genes in both tissue comparisons (Fisher’s exact test *P* < 0.0001, *P* = 0.0002; pollen/leaf and pollen/flower bud respectively). These findings suggest that *R. hastatulus* XY ancestors were mildly pollen biased and *R. rothschildianus* XY ancestors were highly pollen biased, a difference that may be explained by the large difference in levels of Y chromosome degeneration between the species. In particular, because the relatively intact Y chromosomes of *R. hastatulus* still contain a considerable number of genes not expressed in the haploid phase, the bias towards pollen expression should be less severe. Overall, our results suggest that pollen-biased genes may have been involved early in the evolution of sex chromosomes and points to a role for adaptive linkage of haploid beneficial alleles.

To determine whether pollen overexpression evolved on XY-linked gene ancestors after linkage to the sex chromosomes, we performed reciprocal pairwise comparisons of expression patterns of XY genes and their non-XY linked orthologs. Direct comparisons between these gene sets are difficult to interpret due to apparent genome-wide divergence in expression patterns between *R. hastatulus and R. rothschildianus* (Fig. 2). To account for this divergence, we normalized the expression bias of XY and orthologous genes by the average expression bias of their respective autosomal genes to uncover relative differences in expression between species (for details see methods).

We found that XY-linked genes in *R. hastatulus* were significantly more pollen biased than their autosomal orthologs in *R. rothschildianus* in both tissue comparisons (Wilcoxon’s sign rank test *P* < 0.01 both tissue comparisons, Z = −13.58 pollen/leaf, Z = −3.941 pollen/flower bud) (Fig. 3). Similarly, XY-linked genes in *R. rothschildianus* were significantly more pollen biased than *R. hastatulus* orthologs when comparing pollen and flower buds (Wilcoxon’s sign rank test *P* = 0.04, Z = −2.022) but not when comparing pollen and leaf where XY genes appeared to be more pollen biased ancestrally (Wilcoxon’s sign rank test *P* = 0.01, Z = −6.418). Our results support the hypothesis that XY-linked genes play an important role in the haploid gametophytic phase by becoming enriched for pollen expression during (adaptive linkage) and/or following (pollenization) sex chromosome linkage, particularly for *R. hastatulus*. We posit that the lack of pollen expression enrichment observed when comparing leaf and pollen in *R. rothschildianus* may be related to the highly degenerate nature of the Y chromosomes in this species. Long periods of inefficient selection may have eroded signatures of pollenization and/or adaptive linkage.

**Fig. 3:**
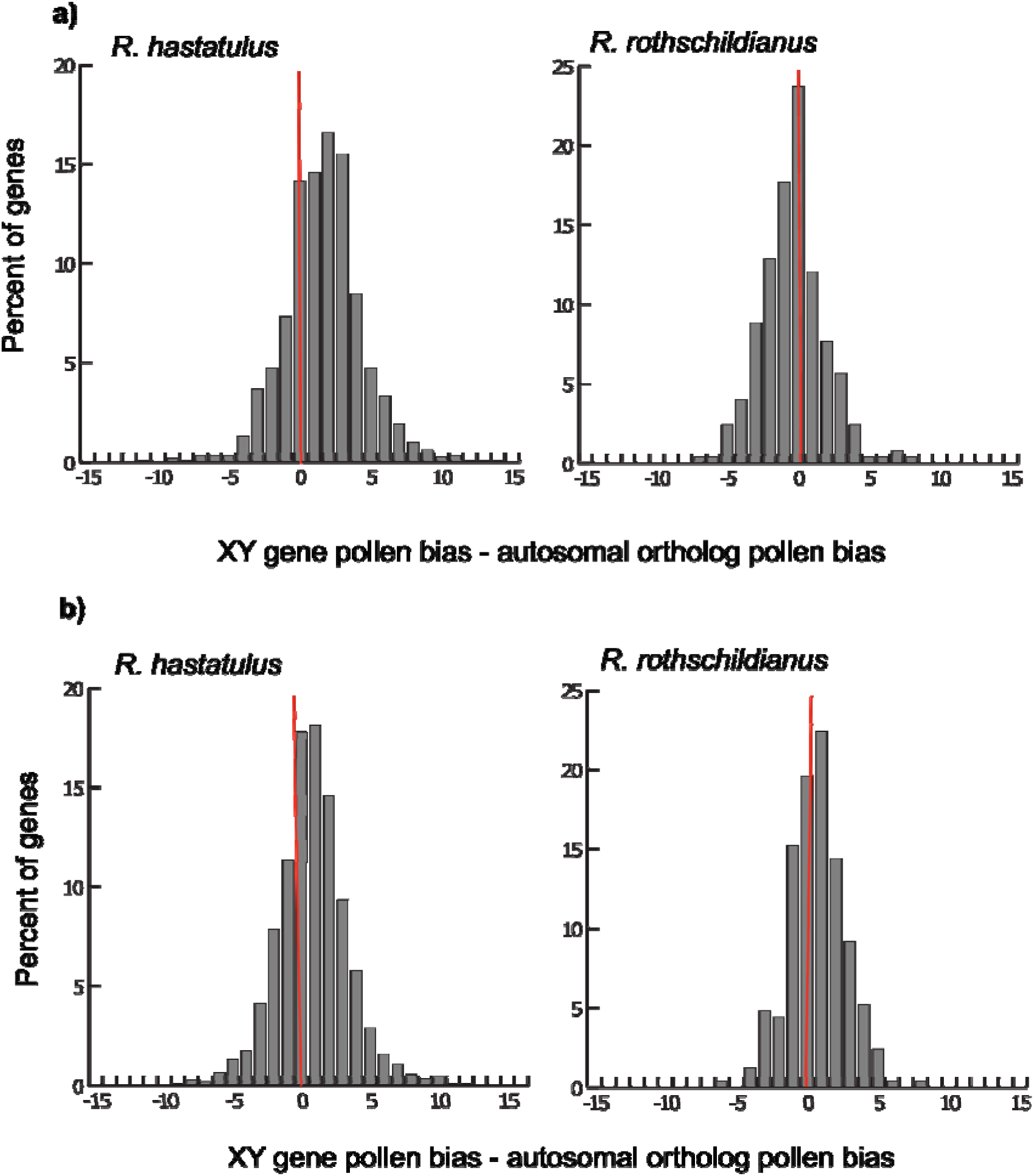
Differences in normalized tissue expression bias of XY genes and their autosomal orthologs in two species of *Rumex*. The magnitude and direction of the differences are related to the evolution of tissue expression in XY genes after their linkage to the sex chromosomes. Tissue expression data used in the comparisons include a) leaf/pollen and b) flower bud/pollen. Positive values indicate greater pollen overexpression in XY genes relative to their autosomal orthologs. For details of normalization see methods.

### Widespread Y overexpression is present specifically in pollen

If pollen overexpression on sex chromosomes is due to adaptive linkage or pollenization on the Y chromosome (Fig. 1a,b), we would predict that it is driven by upregulation of Y-linked genes expressed in pollen. To test this hypothesis, we examined allele-specific gene expression on the sex chromosomes across several tissues. We predicted Y-bias in pollen if adaptive linkage and/or pollenization contribute to the evolution of plant sex chromosomes, and Y-bias in male flower buds if sexual antagonism is the dominant force driving sex chromosome evolution. In contrast, in leaf we predicted minimal Y-bias, or reduced expression on the Y due to degeneration (Hough et al. 2014; Crowson et al. 2017), and therefore we used this tissue as a control.

We found no consistent chromosomal bias for allelic overexpression in leaf (14.6% X overexpressed, 16.4% Y overexpressed) or flower bud (7.7% X overexpressed, 7.0% Y overexpressed) tissue of *R. hastatulus*. There was also no significant difference in the pattern of allelic overexpression between leaf and flower bud tissue (Fisher’s exact test *P* = 0.7083) (Fig. 4). In pollen, however, 44.9% of XY genes exhibited Y overexpression whereas only 16.4% had X overexpression, indicating that XY genes had significantly more Y overexpression in pollen than leaf tissue (Fisher’s exact test *P* = 0.0021).

**Fig. 4:**
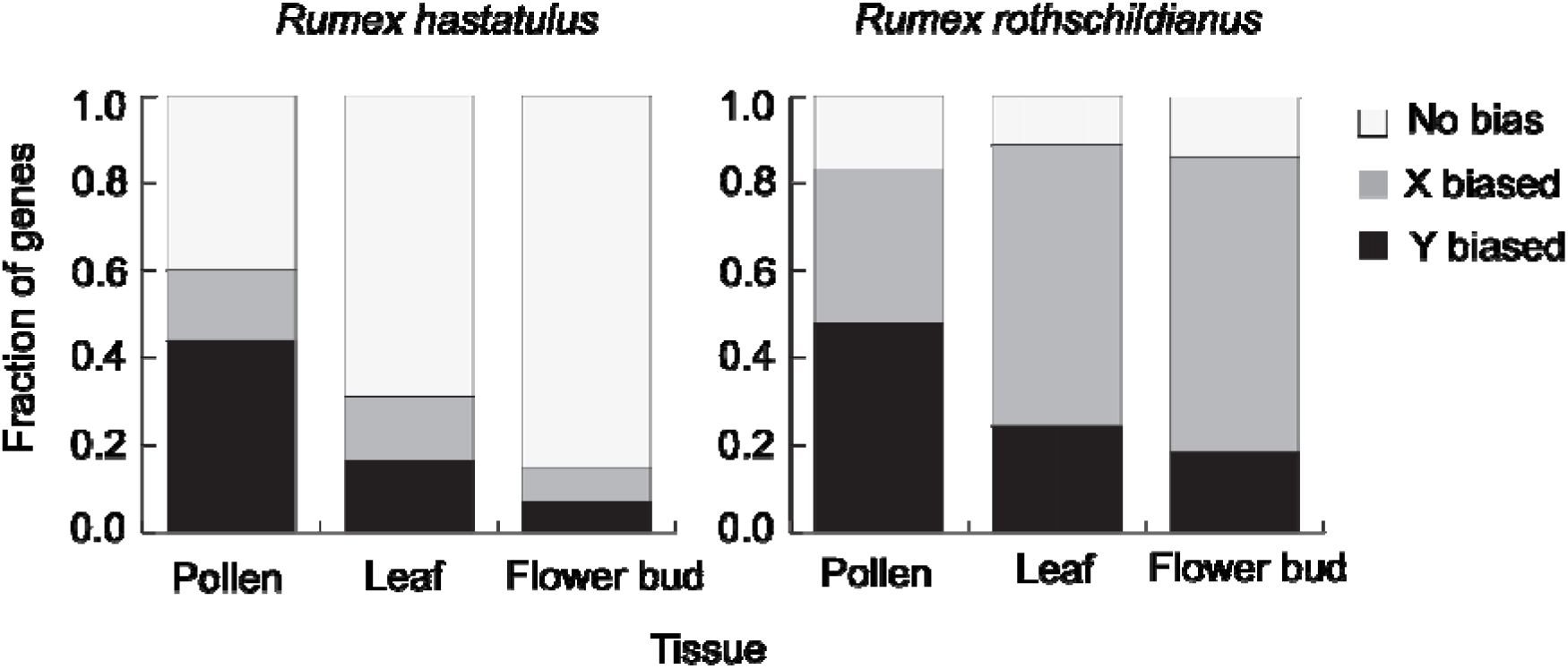
Allele specific expression bias of X- and Y-linked genes in two species of *Rumex*. Bar segments represent the percent of XY genes with no allelic bias (white), significant X-overexpression (grey), and significant Y-overexpression (black).

In *R. rothschildianus* we found overall more X allele-biased genes in flower bud (67.5% X overexpression, 18.2% Y overexpression) and leaf tissue (64.4% X overexpression, 24.3% Y overexpression), with no significant difference in the direction of allele-specific expression between the tissues (Fisher’s exact test, *P* = 0.1167). Once again, we found significantly more Y-biased expression in pollen (47.8% Y overexpressed, 35.5% X overexpressed) relative to leaf tissue (Fisher’s exact test, *P* < 0.0001).

The occurrence of widespread Y-overexpression in pollen of both species is consistent with the hypothesis that sex-specific haploid gametophytic selection plays a significant role in the evolution of sex chromosomes (pollenization and/or adaptive linkage) and suggests that Y-linked alleles are preferentially upregulated in the haploid phase, for which they have been optimised. The overall prevalence of X-overexpression in *R. rothschildianus* confirms previous findings that X-linked alleles appear to be more highly expressed when their Y-linked orthologs accumulate deleterious mutations due to inefficient selection (Crowson et al. 2017). Given that the sex chromosomes of *R. rothschildianus* are far more degenerate than those of *R. hastatulus* it is expected that this pattern of widespread X overexpression is more prominent in *R. rothschildianus*.

We interpret the lack of widespread Y-overexpression in male flower buds as an indicator that sexual antagonism in diploid flower buds contributes less to the evolution of sex chromosomes in the two *Rumex* species than sex-specific haploid selection. Alternatively, it could be that such sexual antagonism is resolved through mechanisms that do not leave a signature in allelic expression patterns. However, this may be unlikely given that sexual antagonism has previously been linked with allele-specific expression changes in plants (Zemp et al. 2016).

Note that given our data it is difficult to definitively disentangle the possible effects of pollenization and adaptive linkage on Y chromosome evolution. We cannot differentiate between a scenario where alleles that are overexpressed in the haploid phase are recruited to the Y (via adaptive linkage), or a scenario where haploid beneficial allele are further upregulated in the haploid phase post linkage (pollenization). Both scenarios are, however, likely complementary and point to the importance of sex-specific haploid selection during plant Y chromosome evolution.

Given the evidence for pollen specialization on the Y chromosome, the early stages of sex chromosome evolution in *Rumex* may have been driven by haploid selection. It has been proposed that this should lead to male-biased sex ratios in populations (Scott and Otto 2017). But contemporary populations of several *Rumex* species, including both species studied here typically show female biased sex ratios (Putwain and Harper 1972; Zarzycki and Rychlewski 1972; Klimes 1993; Rottenberg 1998; Stehlik and Barrett 2005; Pickup and Barrett 2013). This discrepancy may be resolved if we consider the time dependent nature of Y chromosome degeneration (Bachtrog, 2008) and how this might affect sex-ratio evolution.

In particular, early on in Y chromosome evolution male-biased sex ratios may occur because Y degeneration is limited and at an early stage. However, in the long-term, linked selection is likely to cause the accumulation of slightly deleterious mutations on the Y. Given evidence for a very severe loss of diversity on the contemporary Y chromosomes of *R. hastatulus* (Hough et al 2017), the reduced efficacy of selection may be severe enough to drive down haploid fitness on the Y, leading to female-biased sex ratios over time. This view of sex ratio evolution is consistent with comparative data which indicates that plant species with homomorphic sex chromosomes tend to have male-biased sex ratios whereas those with heteromorphic sex chromosomes exhibit female-biased sex ratios (Field et al. 2013). Furthermore, theoretical work indicates that female-biased sex ratios can be maintained in populations following deleterious mutation accumulation on the Y (Hough et al. 2013). Thus, while explicit modelling of this process has yet to be conducted, empirical and theoretical work does suggest that Y degeneration can lead to female-biased sex ratios in older, heteromorphic sex chromosomes, despite a history of specialization for haploid expression on the Y.

It is important to note that we have only considered the role of male haploid selection and it is possible that similar processes could occur in females. Due to the presence of uniovulate flowers in *Rumex* we suspect that gametic competition should be far more intense in males than females. Nonetheless, processes such as meiotic drive during female meiosis may also have significant effects on sex chromosome evolution in these species and contribute to sex chromosome evolution. Future work examining the role of meiotic drive in the spread of recombination suppression on the sex chromosomes would thus be of interest.

## Conclusion

We report evidence that differential retention of pollen-expressed genes during degeneration, pollenization upon divergence, and/or adaptive linkage of pollen-expressed genes jointly contribute to the enrichment of Y chromosomes for pollen expressed genes in *Rumex*. As previously reported in *Silene latifolia* (Chibalina and Filatov 2011), haploid selection can slow the degeneration of Y chromosomes despite the reduced efficacy of selection predicted to be associated with the loss of recombination (Charlesworth 1991). Similar to increased sex-specific expression on animal sex chromosomes, we find sex-specific transmission produces unique conditions in which the Y chromosome becomes specialized for pollen competition. Though our results do not explicitly demonstrate that haploid selection has played a role in driving the evolution of recombination suppression, our findings are consistent with several of the predictions of the model of sex-chromosome evolution proposed by Scott and Otto (2017), whereby sex-specific haploid selection is a key process driving the evolution of sex chromosomes.

## Acknowledgements

The authors would like to thank Daisy Crowson and Tia Harrison for help with RNA extraction from pollen and flower buds. This research was funded by Natural Sciences and Engineering Research Council of Canada Discovery grants (to S.C.H.B. and S.I.W.) and an EWR Steacie fellowship (to S.I.W.).

## Author Contributions

SIW, SCHB and GS conceived of and designed the study, GS collected the data, GS and FEGB analysed the data, all authors contributed to the writing of the paper.

## Conflict of interest

The authors declare no conflict of interest.

